# QInfoMating: Sexual Selection and Assortative Mating Estimation Software

**DOI:** 10.1101/2023.09.06.556585

**Authors:** A. Carvajal-Rodríguez

## Abstract

**Background:** Sexual selection theory is a multifaceted area of evolutionary research that has profound implications across various disciplines, including population genetics, evolutionary ecology, animal behavior, sociology, and psychology. It explores the mechanisms by which certain traits and behaviors evolve due to mate choice and competition within a species. In the context of this theory, the Jeffreys divergence measure, also known as population stability index, plays a key role in quantifying the information obtained when a deviation from random mating occurs for both discrete and continuous data. Despite the critical importance of understanding mating patterns in the context of sexual selection, there is currently no software available that can perform model selection and multimodel inference with quantitative mating data to test hypotheses about the dynamics underlying observed mating patterns. Recognizing this gap, I have developed QInfoMating which provides a comprehensive solution for analyzing and interpreting mating data within the framework of sexual selection theory.

**Results:** The program QInfoMating incorporates a user-friendly interface for performing statistical tests, best-fit model selection, and parameter estimation using multimodel inference for both discrete and continuous mating data. A use case is presented with real data of the species *Echinolittorina malaccana*.

**Conclusions:** The application of information theory, model selection, and parameter estimation using multimodel inference are presented as powerful tools for the analysis of mating data, whether quantitative or categorical. The QInfoMating program is a tool designed to perform this type of analysis.

## Introduction

By this point, there is no doubt about the importance of sexual selection theory and its impact in areas such as population genetics, evolutionary ecology, animal behavior, and even sociology and psychology. There are several strategies for measuring the impact of sexual selection, depending on the field of study and whether the available data pertain to discrete or continuous traits [1–3]. In previous works [2–4], it has been demonstrated that the Jeffreys divergence (also symmetric Kullback-Leibler divergence or population stability index), noted as *J*_PTI_ in [3] corresponds to the increase in information when mating is not random. What this means is that the information measured by the *J_PTI_* statistic is zero if the mating pattern is random, i.e., the mating frequencies match those expected from population frequencies, but *J_PTI_* is greater than zero otherwise.

It is worth clarifying here that in the previously mentioned works as well as in this work, sexual selection and assortative mating are considered patterns reflected in mating data. That is, sexual selection, defined as any type of selection arising from differential fitness in regard to access to gametes for fertilization [5], will produce changes in mating frequencies relative to population frequencies (for any trait involved in it). Assortative mating corresponds to a pattern of matings that are more frequent between similar phenotypes than expected by chance (positive assortative mating) or less frequent than expected by chance (negative assortative mating). Within this framework, the pattern of sexual selection could be produced by two biological processes involving non-random mating: competition to mate and mate choice (see [2] and the references therein).

The process of competition to mate refers, in a broad sense, to access to mating through courtship, intrasexual agression, competition for limited reproductive resources and post-copulatory mechanisms as sperm competition [6–8]. Mate competition may generate a pattern of sexual selection in the sex that competes. The process of mate choice refers to the effects of certain traits expressed in one sex that leads to the non-random allocation of reproductive effort by members of the opposite sex and may be mediated by phenotypic (sensorial or behavioral) properties [9–11]. Mate choice may generate a pattern of assortative mating (positive or negative) and also a pattern of sexual selection [3, 4].

It can be shown [2, 12] that the *J*_PTI_ divergence can be broken down additively into one component that measures the pattern of sexual selection and another that measures assortative mating, and these statistics can be applied both to discrete and continuous data (see below).

The program QInfoMating is a major update (see below) to the Infomating program [3, 4] and incorporates the new quantitative methods developed in [2]. The previous version (Infomating version ≤ 0.4) have already been utilized in empirical studies [13–15].

The methodology incorporated in QInfoMating, which includes both the previously developed approach for discrete characters and the new one for continuous characters, is valid for detecting sexual selection and assortative mating, as well as estimating the best-fitting models of mate choice and mate competition. The goal of the model estimation methodology is to identify the simplest model of mating interactions among phenotypic classes of each sex that can explain observed mating patterns. The modeling framework developed in [4] requires that the data be structured into phenotypic (discrete) classes. Once the data are classified into classes, the model estimation can be applied to any species where correlations and/or frequency tables can be measured in matings for any character of interest

For example, this methodology has been used to study the relationship between color polymorphism and assortative mating in the beetle *Oreina gloriosa* [15], and *Littorina fabalis* [13]. Furthermore, mate choice and shell color polymorphism have been examined in *Littorina saxatilis* [16], and the mechanism of size-based mate choice has been investigated in *Echinolittorina malaccana* [14]. Furthermore, body size similarity has been studied in gastropods in general [17].

While there are other software options available for computing sexual selection and assortative mating from mating tables, such as JMating [18], to the best of my knowledge, QInfoMating (previously Infomating) is the only software for applying model selection theory to mating data, enabling the estimation of sexual selection and assortative mating for both discrete and quantitative data.

## Implementation

The backend of the program is implemented in C++11, while the graphical user interface is implemented in Python 3. Therefore, QInfomating can be run via the user interface (Figure 1-A) or directly from the console under Windows, Linux and macOS. Detailed instructions for use are provided in the program manual (https://acraaj.webs.uvigo.es/InfoMating/QInfoMating_v1.2_Manual.pdf), specifying input formats, the meaning of parameters, models, and result files. In summary, QInfoMating offers the capability to analyze both continuous and discrete data for sexual selection and assortative mating. It begins by performing relevant statistical tests to detect sexual selection and assortative mating and then identifies the best-fitting model. The program employs multi-model inference techniques to estimate parameter values.

**Figure 1.**
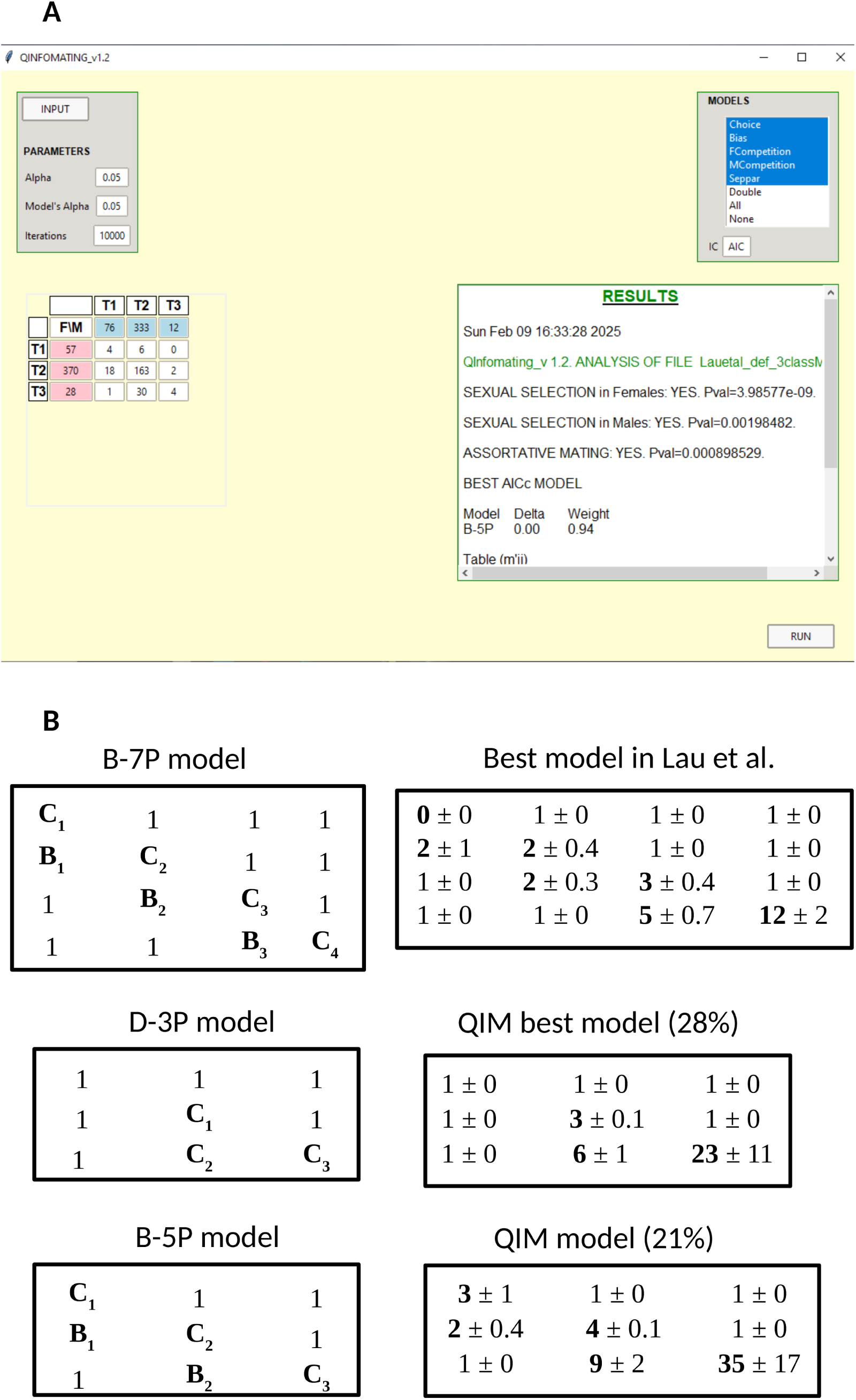
A: QInfomating user-friendly interface. B: Best-fitting models in previous work and in the current analysis. The cells in the tables are parameters of mutual fitness for mating, rows are females and columns are males, so the leftmost cell in the first row is the mutual fitness for mating between the female of class 1 and the male of class 1 and the rightmost cell in the first row is for mating between the female of class 1 and the male of the last class (see text).

### Difference of QInfoMating with the previous version (Infomating)

Unlike the previous version (Infomating), which lacked a user-friendly interface and only analyzed discrete data, QInfomating can be run via the graphical user interface (Figure 1-A) and now accepts continuous data as input (see the format specification in the program manual). When continuous data is provided, the new statistical tests for sexual selection and assortative mating detection, as developed in [2], are performed.

In short, QInfomating replaces Infomating, incorporates a user-friendly interface, and performs all the tasks that Infomating did for discrete traits. Additionally, it includes the new statistical methods developed in [2] to detect patterns of sexual selection and assortative mating for continuous traits and allows for the automatic discretization of these continuous data. This enables model selection analysis regardless of whether the input data is discrete or continuous.

#### Statistical tests

We have already mentioned that the *J_PTI_* divergence can be decomposed additively into a component that measures the pattern of sexual selection and another that measures assortative mating, such that *J_PTI_* = *J_S1_*+ *J_S2_* + *J_PSS_* + *E*, where *J_S1_*and *J_S2_* measure the pattern of sexual selection in females and males, respectively, *J_PSS_* measures the pattern of assortative mating, and *E* is an interaction factor that arises when both patterns are present and that usually has a value close to zero [2].

All *J* components are Jeffreys divergences, what varies are the distributions being compared. In the general case, *J_PTI_* compares the distribution of pairings with the one expected by chance using population data. The *J_S_*_1_ component compares the distribution of females in pairings with the distribution of females in the population, *J_S_*_2_ does the same for males, and *J_PSI_* compares the observed distribution of pairings with the expected random distribution using only the data from the sample of pairs (see [2, 3] for mathematical details). The theoretical formulas for the divergences are the same whether the data is discrete or continuous, but the closed-form formulas for the corresponding statistics change. For example, in the continuous case and assuming a normal distribution, *J_PSI_* can be expressed in terms of correlation (see below), which does not occur in the discrete case.

#### Sexual selection tests

The statistic *J_S1_* compares the distribution of the trait in females that mate with the distribution in the total female population. The pattern of sexual selection in females is detected when these distributions do not match. Similarly, *J_S2_* compares the distribution of the trait in males that mate with the distribution in the total male population (see ref. [2] for a more detailed explanation).

Briefly, we assume that traits *X* and *Y* are normally distributed. The marginal density function for the female trait among females that mate is *f*_1_(x) ∼ *N*(µ_1_, σ_1_^2^), and the trait distribution of the female trait in the entire population is *f*(*x*) ∼ *N*(µ_x_, σ_x_^2^). Then, the test for sexual selection in females is

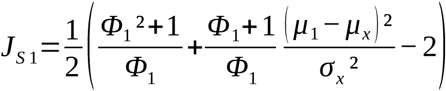

where *Φ*_1_=*σ*_1_^2^/ *σ _x_* ^2^.

And similarly, the marginal density function for the trait in males within matings is *f*_2_(*y*) ∼ *N*(µ_2_, σ_2_^2^) and in the population is *g*(*y*) ∼ *N*(µ_y_, σ_y_^2^) then, the test for sexual selection in males is

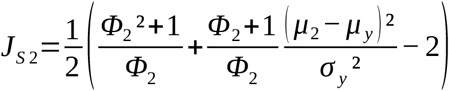

where *Φ*_2_=*σ*_2_^2^/ *σ _y_* ^2^

and *J_PSS_*= *J_S_*_1_+*J_S_*_2_.

For a random sample of *n* matings, *nJ*_S1_ and *nJ*_S2_ are asymptotically χ^2^ distributed with 2 degrees of freedom [19, 20] under the null hypothesis that the females (males) within matings are distributed with mean µ_x_ (µ_y_) and variance σ_x_^2^ (σ_y_^2^) i.e. absence of sexual selection.

If we want to compare only the mean value of the female trait *X* in the mating sample with that of the total female population, we set Φ_1_=1 so that

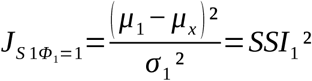

where *SSI*_1_ is the standardized sexual selection intensity index [21–23].

And similarly for males

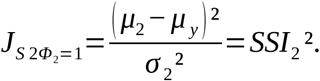

Then with a sample size of *n* matings we have (*nJ*_S1Φ=1_)^0.5^ ∼ *t_n_*_-1_ and (*nJ*_S2Φ=1_)^0.5^ ∼ *t_n_*_-1_.

For large sample sizes, both types of tests (χ^2^ or *t*) work similarly. However, for smaller sample sizes, the χ^2^ test might be slightly liberal. More generally, the χ^2^ test offers a multifaceted approach as it can also detect differences in variances, such as stabilizing (variance reduction) or diversifying (variance increasing) selection, or combinations of these forms of selection with directional selection (differences in means).

#### Assortative mating test

In the case of the assortative mating pattern, we use only the mating distribution and compare the expected distribution by chance with the observed distribution. The assortative mating pattern is detected when the observed distribution differs significantly from that expected by chance.

It can be shown that, assuming the quantitative traits we are measuring are normally distributed and jointly normally distributed within mating pairs, the information measure of assortative mating, *J_PSI_*, that compares the distributions mentioned above, can be expressed as a function of the square of correlation coefficient ρ of the mating pairs that takes values within the set of non-negative real numbers [0, +∞), specifically (see ref [2] for a more detailed explanation):

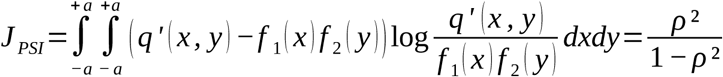

where *q*’(*x*, *y*) is the mating probability density for pairs with values in the infinitesimal interval ([*x*, *x* + *dx*], [*y*, *y* +*dy*]) and *f*_1_(*x*) and *f*_2_(*y*) are as before, the marginal density functions for the female trait among females that mate and for the male trair among males that mate, respectively. For simplicity and without loss of generality we have assumed that the range is the same for *X* and *Y* so that, *X, Y* ∈[*−a, a*] with |*a*| > 0.

For a random sample of *n* matings, *nJ_PSI_* is asymptotically χ^2^ distributed with 1 degree of freedom under the null hypothesis that the mating probability distribution is equal to the product of the female and male marginal distributions within matings [20].

#### Class frequency distribution

Once the statistical tests have been carried out, model selection is only executed if any of the statistical tests yield a significant result. Following this, the program automatically calculates the frequency distribution (classes) for the continuous data and proceeds with multi-model estimation. The algorithm used to obtain the frequency distribution is detailed in the program manual. It is a procedure for discretizing continuous data into bins, which essentially consists of:

1. The range of the data set is calculated (maximum value *Max* - minimum value *Min*).
2. The formula that relates number of classes *K* and sample size *n* is [24]

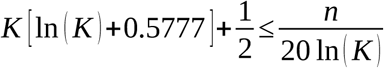

where we conservatively assume a sample size more than one order of magnitude larger than that required for the uniform frequency case (details in the program manual).
3. The width *w* of each class is computed as *w* = (range / *K*).
4. Make the classes. The first class is always [*Min*, *Min* + w), the following classes are [*x*, *x* + *w*) where *x* is the maximum of the range of the previous class and the last class is [*Max* - *w*, *Max*].
5. Calculate the frequency of each class. Check that it is represented in each sex; if not, reduce *K* by one and repeat from 3.

#### Set of models

The set of models used by the program is based on the concept of mutual mating fitness [2, 12] and was developed in previous work [4]. Comparing different mutual mating fitnesses between discrete phenotypic classes allows distinguishing between different models. After the program’s execution, an output file is generated, which includes an explanatory summary of the models used. I will now outline the main differences between these models.

The null model, called *M*_0_, corresponds to random mating, where all mating fitnesses are equal. The most complex model is the saturated model, *M*_sat_, which assumes that all pair of classes have different mutual mating fitness.

Between these two extremes, there are competition models that produce patterns of sexual selection in males and/or females. The names of these models start with

- Sfem- if they produce sexual selection only in females,
- Smale- if only in males,
- S2- if in both.

For example, males of a phenotypic class may systematically have higher mating fitness than other males and this scenario will be captured in Smale-models.

Additionally, there are models that incorporate mate choice and produce an assortative mating pattern. In some cases, these can also result in sexual selection in males, females, or both. Models that generate patterns of sexual selection and assortative mating are denoted in various ways, depending on the parameters involved (see Table 1 in [4]).

**Table 1.** Mating table for the Wild data used in Lau et al[14]. The data is classified in four classes.

| Premating males numbers (421) |  |  |  |
| --- | --- | --- | --- |
| 23 | 181 | 167 | 50 |
| Premating female numbers (455) |  |  |  |
| 23 | 146 | 190 | 96 |
| Matings (228, rows are females) |  |  |  |
| 0 | 2 | 0 | 0 |
| 3 | 30 | 14 | 1 |
| 2 | 42 | 45 | 9 |
| 1 | 8 | 43 | 28 |

Mate choice models that only produce an assortative mating pattern (without sexual selection, as they assume certain conditions in phenotypic frequencies; see Figure 1 and Equation 5 in [4]) always start with C- and imply a preference for similar or dissimilar mates. A variant of these models are the bias models, whose names start with B- and always imply a preference for similar mates but with a bias, for example, a preference for someone slightly larger than oneself. The bias models can also produce sexual selection pattern.

There are also differences in the set of models considered for discrete and continuous cases. In the discrete case, there is a default set of models that can be modified by the user. Conversely, for the continuous case, the program, rather than the user, selects models based on the statistical significance of various tests. This is because, in the continuous case, data are only discretized if the continuous tests are significant. Additionally, the mating models framework developed in [4] applies only to discrete traits. Therefore, model selection techniques will only be applied if the data are discretized. Consequently, with continuous data, we do not know a priori if the model estimates will be performed. Thus, it is more practical for the program to automatically generate only those sets of models, always including the null (random mating) and the saturated, that make sense to explore.

Specifically, if there is a statistically significant pattern of assortative mating but no sexual selection, the program uses the set of models that incorporate mate choice and bias in conjunction with random mating and the saturated model, but it does not include competition models. On the other hand, if there is a statistically significant pattern of both assortative mating and sexual selection in both sexes, QInfoMating includes also models that generate both effects (called double-effects models) among the models to be tested.

In any case, if we prefer to choose the models ourselves, we can do so using the mating table that QInfoMating generates from the quantitative data, which allows us to create an input file for discrete analysis, using the default set of models or any of our choice.

As in the previous version, the program selects the best model according to the selected criterion (Akaike information criterion, AIC, by default). Additionally, it performs multimodel-based inference to estimate the parameters of interest, using a group of models rather than relying on a single best-fit model [4, 25].

## Results

### Real data use case

*Echinolittorina malaccana*, a marine intertidal snail commonly found on rocky shores in the Indo-West Pacific region [26], exhibits mate selection behaviors. Males have shown a clear preference for females slightly larger than themselves [22]. In a subsequent study conducted by [14], the continuous trait of size (specifically, shell length) was transformed into categorical variables by creating four distinct size classes. This transformation allowed for the creation of a mating table (see Table 1), which enabled the utilization of the multimodel inference techniques available in the previous version of the program (Infomating). This approach provided a more profound insight into the intricate dynamics of mate choice. The analysis ultimately confirmed that the most suitable model for mate choice in *Echinolittorina malaccana* involves a size bias, resulting in both assortative mating and the presence of sexual selection favoring larger individuals.

In a recent study [2], the same onshore data, originally presented in [14], and accessible via [27], was directly used without converting the continuous shell length trait into discrete classes. This approach aimed to demonstrate the computation of Jeffreys divergence indices [3] for quantitative traits. The research revealed that the statistical tests for the quantitative data yielded similar conclusions as the previous analysis using the statistical tests for discrete data. However, the quantitative data lacked the necessary structure for applying the model selection methodology.

In this current study, we will leverage QInfoMating to conduct a quantitative analysis of the same data set. The program will apply the quantitative statistical tests and automatically transform data into classes to facilitate model selection and multi-model inference. This analysis aims to compare the results with those obtained in [14]. For consistency, we will employ the Akaike Information Criterion (AIC), the same criterion used in [6], and additionally, the more conservative Bayesian Information Criterion (BIC) to showcase the software’s versatility and utility.

A CSV file containing data on the shell length of each snail in the population (females and males, first two columns) and the phenotype pairs that mated (third and fourth columns) served as input for QInfoMating. Please refer to the input files section of the Program Manual (Figure 3 in the Manual) for the exact input format. Although the original study by Lau et al. in [14], used four classes, QInfoMating segmented the data into three classes (Table 2), before proceeding with the model selection process.

**Table 2.** Mating table generated by QInfoMating for the Wild data in Lau et al [14]. The data is classified in three classes.

| Premating males numbers (421) |  |  |
| --- | --- | --- |
| 76 | 333 | 12 |
| Premating female numbers (455) |  |  |
| 57 | 370 | 28 |

| Matings (228, rows are females) |  |  |
| --- | --- | --- |
| 4 | 6 | 0 |
| 18 | 163 | 2 |
| 1 | 30 | 4 |

The tables shown in Figure 1-B correspond to the models selected by the program. Each cell in the tables represents mutual mating fitness parameters. The rows represent females, and the columns represent males. Therefore, the first cell corresponds to the mating of class 1 females with class 1 males, the second cell in the first row corresponds to class 1 females with class 2 males, and so on.

In the figure, the tables on the left show the model with its parameters, while the tables on the right show the same model with the obtained estimates. First, the best-fitting model by Lau et al. (B-7P, a bias model with 7 parameters) is presented. This is followed by the two best-fitting models obtained by QInfoMating: D-3P (a double-effect model with 3 parameters) and B-5P (a bias model with 5 parameters).

The D-3P model is a model with double effect i.e., generates sexual selection and assortative mating patterns. From Figure 1-B in the D-3P model we appreciate that the choice parameters (main diagonal) are c_1_ and c_3_. The parameter c_1_ can also be considered a male sexual selection parameter jointly with c_2_ since they change the marginal propensity for the second class males (second column). However, from the point of view of biased choice, the parameter c_2_ can be seen as a bias (subdiagonal) in a model in which individuals in the shorter class have more difficulty mating. In fact, the second best model was B-5P, which is the equivalent model to B-7P when there are three classes instead of four.

The best model by Lau et al. had a weight of 99%, so it was almost exact to the multi-model inference parameter matrix. In the case of QInfoMating, the best model (D-3P) had only 28% of the weight and other models such as B-5P (21%), D-5P (21%), D-4P (18%) and D-6P (8%), are necessary to capture more than 95% of the weight. The pattern obtained using the Bayesian Information Criterion (BIC) was similar to that obtained using the AIC criterion but with greater weight concentrated in the D-3P model (Table 3).

**Table 3.** Models that accumulate up to 95% of the weight. The parameter notation corresponds to the B-7P model in Figure 1B where C_i_ represent the ith cell on main diagonal and B_i_ on the subdiagonal.

| Analysis | Models (>95%) | Weight | Parameter estimation |  |  |  |  |  |  |
| --- | --- | --- | --- | --- | --- | --- | --- | --- | --- |
|  |  |  | C<sub>1</sub> | C<sub>2</sub> | C<sub>3</sub> | C<sub>4</sub> | B<sub>1</sub> | B<sub>2</sub> | B<sub>3</sub> |
| Lau et al 2021 | B-7P | 99% | 0 | 2 | 3 | 12 | 2 | 2 | 5 |
| QInfoMating-AIC | D-3P | 28% | 1 | 3 | 22 | - | 1 | 6 | - |
|  | B-5P | 21% | 3 | 4 | 35 | - | 2 | 9 | - |
|  | D-5P | 21% | 3 | 4 | 35 | - | 2 | 9 | - |
|  | D-4P | 18% | 2 | 3 | 24 | - | 1 | 6 | - |
|  | D-6P | 8% | 3 | 4 | 36 | - | 2 | 10 | - |
| QInfoMating-BIC | D-3P | 85% | 1 | 3 | 23 | - | 1 | 6 | - |
|  | D-4P | 10% | 2 | 3 | 24 | - | 1 | 6 | - |
|  | B-5P | 2% | 3 | 4 | 35 | - | 2 | 9 | - |
|  | D-5P | 2% | 3 | 4 | 35 | - | 2 | 9 | - |

However, the structure of all these models is very similar: large males (rightmost columns) choose females larger (closest to the bottom rows) than themselves, especially for the largest classes (B_2_ and B_3_ parameters in the bias models, or c_2_, c_3_ in the D-3P model).

The pattern behind the QInfoMating models aligns with the findings reported by Lau et al. [14] for the Multiple-design and Wild data, as depicted in their Figure 4. However, there is a notable distinction: Lau et al. observed that the shorter class exhibited considerably lower mating ability. This difference may be attributed to the presence of their first class with a minimal number of individuals for both females and males, as documented in Table 1.

However, one might wonder why QInfoMating did not clearly favor the B-5P model, which is most akin to the B-7P model chosen in the original analysis by Lau et al [14]. The explanation lies in how QInfoMating operates when handling quantitative data. As explained before, the program selects models based on the statistical significance of various tests and included double-effect models (as D-3P, D-4P and D-6P) among the set of models to be tested. These particular models were not considered in the original Lau et al. study.

To further elucidate, I excluded the double-effect models and repeated the analysis using exactly the same set of models as in Lau et al. Upon conducting this analysis, the best-fitting model becomes the B-5P, with a weight of 94%. The program’s preference for the D-3P model, when included, stems from its ability to explain the data with two fewer parameters, thus aligning with the principle of parsimony. In cases where both models are closely weighted, the choice between them is influenced by the researcher’s experience and the biological plausibility of the results. Nonetheless, it’s worth noting that the fundamental structure of these models remains quite similar: larger males tend to prefer females larger than themselves, whenever such a pairing is possible.

Consistent with the result of models, the statistical tests performed by QInfoMating detect assortative mating and sexual selection both in females and males (Table 4).

**Table 4.** QInfoMating tests for the Wild data used in Lau et al [14]. For the discrete tests the data is classified in three classes. *J_PSI_* is the assortative mating test. *J_S_*_1_ and *J_S_*_2_ are the sexual selection tests for females and males, respectively.

| Test type | Statistic | $H_0$ | $p$-value |
| --- | --- | --- | --- |
| Quantitative | $J_{PSI} = 0.49$ | no assortative mating | $<10^{-6}$ |
| Quantitative | $J_{S1} = 0.24$ | no female sexual selection | $<10^{-6}$ |
| Quantitative | $J_{S2} = 0.07$ | no male sexual selection | 0.0004 |
| Discrete | $J_{PSI} = 0.08$ | no assortative mating | 0.0009 |
| Discrete | $J_{S1} = 0.17$ | no female sexual selection | $<10^{-6}$ |
| Discrete | $J_{S2} = 0.06$ | no male sexual selection | 0.002 |

## Discussion

Assortative mating based on size and other quantitative traits is common in many species, from gastropods [22] to humans [28]. The processes leading to assortative mating may not solely rely on similarity-based mechanisms and may not affect different phenotypic classes equally. This is where the power of model selection methodology becomes evident, as it doesn’t require proposing a single ad hoc model. Instead, it allows best fit of data for a wide range of models, enabling researchers to study non-trivial mating patterns, some of which may not have been obvious before the experiment.

## Conclusion

In conclusion, the application of information theory, model selection, and parameter estimation using multimodel inference are presented as powerful tools for the analysis of mating data, whether quantitative or categorical. QInfoMating is software designed to perform this type of analysis.

## Availability and requirements

**Project name:** QInfomating

**Project home pages:** https://noosdev0.github.io/QInfoMating

also

https://acraaj.webs.uvigo.es/InfoMating/QInfomating.htm

**Operating system(s):** Windows, Linux, macOS

**Programming language:** C++, Python.

**License:** GNU GPL.

## Declarations

## Data availability

The datasets generated and/or analysed during the current study are available in the DRYAD repository, http://doi.org/10.5061/dryad.h214h8t

## Competing interests

The authors declare that they have no competing interests

## Funding

This was funded by Xunta de Galicia (ED431C 2024/22), Ministerio de Ciencia e Innovación (PID2022-137935NB-I00) and Centro singular de investigación de Galicia accreditation 2024-2027 (ED431G 2023/07) and “ERDF A way of making Europe”. Funding for open access charge: Universidade de Vigo/ CRUE-CISUG.

## Authors’ contributions

ACR is the sole author of this work.

## Acknowledgments

Dedicated to my sister Paci. I thank E. Rolán-Alvarez for comments on a previous version of the manuscript.

## Notes

### Competing Interest Statement

The authors have declared no competing interest.

### Summary of Updates

Improved models explanation and figure and table captions. Other minor text corrections.

## References

1. Henshaw JM, Kahn AT, Fritzsche K. A rigorous comparison of sexual selection indexes via simulations of diverse mating systems. Proc Natl Acad Sci. 2016;113:E300–8.

2. Carvajal-Rodríguez A. Unifying quantification methods for sexual selection and assortative mating using information theory. Theor Popul Biol. 2024;158:206–15.

3. Carvajal-Rodríguez A. Non-random mating and information theory. Theor Popul Biol. 2018;120:103–13.

4. Carvajal-Rodríguez A. Multi-model inference of non-random mating from an information theoretic approach. Theor Popul Biol. 2020;131:38–53.

5. Shuker DM, Kvarnemo C. The definition of sexual selection. Behav Ecol. 2021;32:781–94.

6. Andersson M. Sexual selection. Princeton, N.J.: Princeton University Press; 1994.

7. Kokko H, Klug H, Jennions MD. Unifying cornerstones of sexual selection: operational sex ratio, Bateman gradient and the scope for competitive investment. Ecol Lett. 2012;15:1340–51.

8. Wacker S, Amundsen T. Mate competition and resource competition are inter related in sexual selection. J Evol Biol. 2014;27:466–77.

9. Edward DA. The many facets of mate choice: a response to comments on Edward. Behav Ecol. 2015. 10.1093/beheco/arv021.

10. Rosenthal GG. Mate choice: the evolution of sexual decision making from microbes to humans. Princeton University Press; 2017.

11. Jennions MD, Petrie M. Variation in mate choice and mating preferences: a review of causes and consequences. Biol Rev. 1997;72:283.

12. Carvajal-Rodríguez A. On Non-Random Mating, Adaptive Evolution, and Information Theory. Biology. 2024;13:970.

13. Estévez D, Kozminsky E, Carvajal-Rodríguez A, Caballero A, Faria R, Galindo J, et al. Mate Choice Contributes to the Maintenance of Shell Color Polymorphism in a Marine Snail via Frequency-Dependent Sexual Selection. Front Mar Sci. 2020;7.

14. Lau SLY, Williams GA, Carvajal-Rodríguez A, Rolán-Alvarez E. An integrated approach to infer the mechanisms of mate choice for size. Anim Behav. 2021;175:33–43.

15. Roggero A, Alù D, Laini A, Rolando A, Palestrini C. Color polymorphism and mating trends in a population of the alpine leaf beetle Oreina gloriosa. Plos One. 2024;19:e0298330.

16. Gefaell J, Vigo R, Galindo J, Rolán-Alvarez E. Experimental evidence of mate choice as the driving mechanism behind negative assortative mating for shell colour in a marine snail. Biol J Linn Soc. 2024;142:441–51.

17. López-Cortegano E, Carpena-Catoira C, Carvajal-Rodríguez A, Rolán-Alvarez E. Mate choice based on body size similarity in sexually dimorphic populations causes strong sexual selection. Anim Behav. 2020;160:69–78.

18. Carvajal-Rodríguez A, Rolan-Alvarez E. JMATING: a software for the analysis of sexual selection and sexual isolation effects from mating frequency data. BMC Evol Biol. 2006;6:40.

19. Evren A, Tuna E. On some properties of goodness of fit measures based on statistical entropy. Int J Res Rev Appl Sci. 2012;13:192–205.

20. Kullback S. Information Theory and Statistics. New edition. Mineola, N.Y: Dover Publications; 1997.

21. Erlandsson J, Rolán-Alvarez E. Sexual selection and assortative mating by size and their roles in the maintenance of a polymorphism in Swedish *Littorina saxatilis* populations. Hydrobiologia. 1998;378:59–69.

22. Ng TPT, Rolán-Alvarez E, Dahlén SS, Davies MS, Estévez D, Stafford R, et al. The causal relationship between sexual selection and sexual size dimorphism in marine gastropods. Anim Behav. 2019;148:53–62.

23. Caballero A. Quantitative Genetics. Cambridge University Press; 2020.

24. Lewontin RC, Prout T. Estimation of the Number of Different Classes in a Population. Biometrics. 1956;12:211–23.

25. Burnham KP, Anderson DR, Huyvaert KP. AIC model selection and multimodel inference in behavioral ecology: some background, observations, and comparisons. Behav Ecol Sociobiol. 2011;65:23–35.

26. Reid DG. The genus *Echinolittorina* Habe, 1956 (Gastropoda: Littorinidae) in the Indo-West Pacific Ocean. Zootaxa. 2007;1420:1–161.

27. Rolán-Alvarez E, Williams G, Ng T, Estévez-Barcia D, Davies M, Saltin S, et al. Data for: The causal relationship between sexual selection and sexual size dimorphism in marine gastropods. 2021;1.

28. Robinson MR, Kleinman A, Graff M, Vinkhuyzen AAE, Couper D, Miller MB, et al. Genetic evidence of assortative mating in humans. Nat Hum Behav. 2017;1:1–13.

